# Energetics and Conformational Pathways of Functional Rotation in the Multidrug Transporter AcrB

**DOI:** 10.1101/173567

**Authors:** Yasuhiro Matsunaga, Tsutomu Yamane, Tohru Terada, Kei Moritsugu, Hiroshi Fujisaki, Satoshi Murakami, Mitsunori Ikeguchi, Akinori Kidera

## Abstract

The multidrug transporter AcrB transports a broad range of drugs out of the cell by means of the proton-motive force. The asymmetric crystal structure of trimeric AcrB suggests a functionally rotating mechanism for drug transport. Despite various supportive evidences from biochemical and simulation studies for this mechanism, the link between the functional rotation and proton translocation across the membrane remains elusive. Here, calculating the minimum free energy pathway of the functional rotation for the complete AcrB trimer, we describe the structural and energetic basis behind the coupling between the functional rotation and the proton translocation at atomic-level. Free energy calculations show that protonation of Asp408 in the transmembrane portion of the drug-bound protomer drives the functional rotation. The conformational pathway identifies vertical shear motions among several transmembrane helices, which regulates alternate access of water in the transmembrane as well as peristaltic motions pumping drugs in the periplasm.

---

Bacterial multidrug resistance (MDR) is an increasing threat to current antibiotic therapy^1^. Resistance nodulation cell division (RND) transporters are one of the main causes of MDR in Gram-negative bacteria including human pathogens. These transporters pump a wide spectrum of antibiotics out of the cell by means of proton or sodium motive forces, conferring MDR to the bacterium when overexpressed. Understanding the mechanism of the drug efflux process is invaluable for the treatment of bacterial infections and the design of more effective drugs or inhibitors.

In *E*. *coli*, the AcrA-AcrB-TolC complex is largely responsible for MDR against many lipophilic antibiotics^2^. This tripartite assembly spans the periplasmic space between the inner and the outer membranes of the cell, transporting drugs from the cell to the medium, bypassing periplasm and the outer membrane. TolC, an outer membrane protein, forms a generic outer membrane channel in which drugs can passively move towards the medium. AcrB is an inner membrane protein, primarily responsible for specificity towards drugs and their uptake, as well as for energy transduction. AcrA acts as an adaptor which bridges TolC and AcrB^3^.

AcrB is one of the best characterized RND transporters in both experiments and simulations, making it a prototype for studying the drug efflux mechanism of the RND family^4^. AcrB transports drugs from the inner membrane surface or the periplasm to the TolC channel using the proton-motive force. The structure of AcrB was first solved in a threefold symmetric form^5^, and later in an asymmetric form^6-8^. AcrB is a homotrimer with a triangular-prism shape, and each protomer is made up of three domains (Fig. 1a): the transmembrane (TM) domain and an extensive periplasmic portion comprising the porter and the funnel domains. The TM domain transfers protons across the inner membrane down the electrochemical gradient (Fig. 1c). The porter domain is made of four subdomains, PN1, PN2, PC1 and PC2 (Fig. 1b), and is responsible for drug transportation. The funnel domain has a funnel-like shape with an exit pore that indirectly connects to TolC via AcrA^3^.

**Figure 1:**
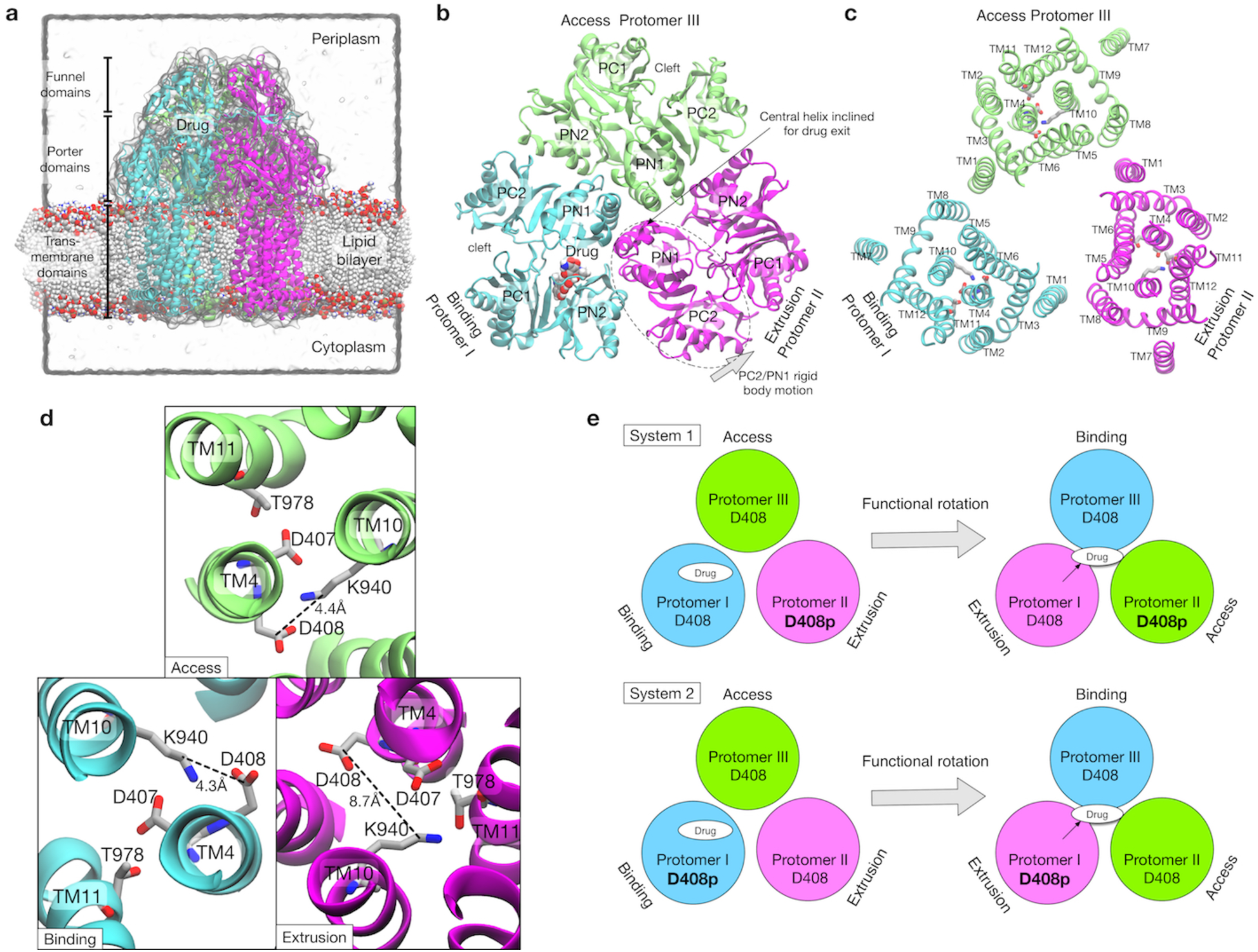
Crystal structure of AcrB and simulation setup. **(a)** Simulation box and a side view of AcrB embedded in POPE lipid bilayer. A drug (minocycline) is represented by spheres. **(b)** Porter domains of the crystal structure viewed from the cell exterior. The protomer in cyan, magenta, lime represents the Binding, the Extrusion, and the Access states, respectively. **(c)** The transmembrane domain viewed from the cell exterior. Key residues related to proton translocation are represented by sticks. **(d)** Close-up view of the key residues. **(e)** Schematics of the simulation systems. The protonated D408 is denoted by D408p. System 1 has D408p in the Protomer II. System 2 has D408p in the Protomer I.

In the asymmetric structure, each protomer of AcrB adopts a distinct conformation which assumes one of the three functional states in the drug transport cycle; Access^6^ (or Loose^4,7^), Binding (Tight) and Extrusion (Open) states, or A, B and E, respectively. The A state has entrances (an open cleft between PC1 and PC2, and a groove between TM helices, TM8 and 9) for drug uptake. The B state has a binding pocket (called the distal binding pocket) formed by PN1, PN2 and PC1 (Fig. 1b). This pocket contains many hydrophobic residues, as well as several charged and polar ones, contributing to broad substrate “polyspecificity”^9,10^. The E state has a drug exit pathway to the exit pore formed by an inclined central helix (Fig. 1b). The tilt of the central helix is achieved by rigid body motions of a PC2/PN1 repeat (see Fig. 1b and Supplementary Fig. S7a). This motion also closes the PC1/PC2 cleft, leading to contraction of the binding pockets^4^.

These findings from the asymmetric structure and other biochemical data led to the proposal of the functional rotation mechanism^6,7^. Hereafter, following Yao et al.^11^, we use a three letter notation to represent the state of the trimer. For example, the BEA state represents a trimeric state where the protomer I is in the B state, the protomer II in the E state, and the protomer III in the A state (where protomers are numbered in a counterclockwise order). In the functional rotation mechanism, for example starting from the BEA state, drug efflux is coupled with a conformational transition to the EAB state. Viewed from the top, this transition is a 120 degrees rotation of functional states (not a physical rotation). This mechanism is supported by both biochemical studies^12,13^, and molecular dynamics (MD) simulation studies^11,14^.

The three functional states satisfy the conditions required in the alternate access model^15^: the A state (and possibly also the B state) corresponds to an inward-facing state, the E state represents an outward-facing state, and the B state is an occluded state. However, despite this excellent correspondence between the structures and states, dynamical picture of functional rotation remains unclear. Specifically, how are the conformational changes (functional rotation) and the energy source (proton translocation) synchronized and coupled dynamically? In contrast to small transporters, where substrate efflux and energy transduction are spatially coupled in the TM domain^16^, drug efflux by AcrB occurs in the periplasmic space, spatially separate (~50 Å apart) from the proton translocation sites in the TM domain. To understand the whole process of drug efflux of AcrB, the energy transduction mechanism between the two distant sites should be clarified, which is the purpose of this study.

The TM domain of each protomer contains twelve α-helices (TM1-12, Fig. 1c). Essential residues for proton translocation are in the middle of the TM helices, D407 and D408 on TM4, K940 on TM10, and R971 and T978 on TM11. AcrB mutants with these residues substituted by alanine are inactive^17,18^. In the asymmetric structure, K940 shows a distinct side chain orientation in the E state compared with that in the A and B states (Fig. 1d). Specifically, K940 tilts away from D408, and towards D407 and T978 in the E state, suggesting that transient protonation of D408 contributes to the TM conformational change. Recently, all-atom MD simulation by Yamane et al. showed that the protonation of D408 stabilized the E state, while deprotonated D408 induced the conformation change of the TM domain toward the A state^19^. Eicher et al. showed by X-ray crystallography of AcrB mutants combined with MD simulations that these conformational changes in the TM domain were related to the alternate access of water, and in turn to the functional rotation.^20^.

Although these studies provide invaluable insights into the energy transduction mechanism, the difficulty in experimental observation of the protonation states and the time-scale limitation in all-atom MD simulations preclude direct characterization of the energy transduction process. The most direct characterization would be the observation of the functional rotation in various protonation states. To achieve this, we conducted extensive MD simulations with state-of-the-art supercomputing^21^. Using an explicit solvent/lipid all-atom model (480,074 atoms), we identified conformational transition pathways in one step of the functional rotation for the complete AcrB trimers with different protonation states. Specifically, the physically most probable conformational pathway (the minimum free energy pathway) was searched by the string method^22-30^, an efficient parallel algorithm which overcomes the time-scale limitation in MD simulations. By comparing energetics and conformational pathways of two systems with each different protonation state, we clarify the relationship between the protonation and the functional rotation. Free energy evaluation along the pathways shows that protonation of D408 in the B state induces the functional rotation. Structural analysis along the conformational pathway reveals that vertical shear motions in specific TM helices regulate the alternate access of water in the TM domain as well as the peristaltic motions pumping a drug in the porter domain. These findings provide a simple and unified view of energy transduction in AcrB.

## Results

### Energetics of functional rotation under two different protonation states

We computed the most probable conformational pathway in the functional rotation from the BEA state to the EAB state with the string method. From the observation of the crystal structure, we postulated that the protonation of D408 in the B state induces the conformational change of toward the E state. This hypothesis was tested with the setup of two different simulation systems (Fig. 1e): System 1 has a protonated D408 (hereafter, denoted by D408p) in protomer II, which undergoes a transition from the E to the A state. System 2 has D408p in protomer I, undergoing from the B to the E state. If the hypothesis is correct, the system 1 should be stable at the initial BEA state while the system 2 prefers the functional rotation. A drug (minocycline) was explicitly included in both systems. It is bound in protomer I (the B state) at first, and then transported to the exit pore during the functional rotation. No additional drug for drug uptake was included in order to keep the drug-AcrB interaction as simple as possible. For the string method calculation, collective variables (Cartesian coordinates of Cα atoms in the porter domain and TM helices, see Methods and Supplementary Fig. S1) were carefully chosen according to previous simulation studies^11,19^. The drug was not included in the collective variables since it is diffusive. The pathway in the collective variable space was discretized by 30 points (called images). Image 1 corresponds to the BEA state, and image 30 is the EAB state.

Figure 2a shows the free energy profiles along the pathways obtained by the string method, the convergence towards the minimum free energy pathway is shown. The impact of protonation is revealed when the converged pathways of the two systems are compared (see Fig. 2b): While system 1 has an energy minimum close to the initial BEA state (at image 5), the minimum of system 2 is shifted towards the final EAB state (at image 15), indicating that the protonation state in system 2 drives the functional rotation. The increase in free energy in the late stage (images 20-30) of both systems is partially caused by the release of drug with no uptake of a new drug molecule, which may be needed to progress to the next step of rotation. Previous simulation studies^11,19^ suggested that asymmetric trimeric states are unstable without a bound drug while the symmetric AAA state becomes most stable, avoiding rebinding of the released drug.

**Figure 2:**
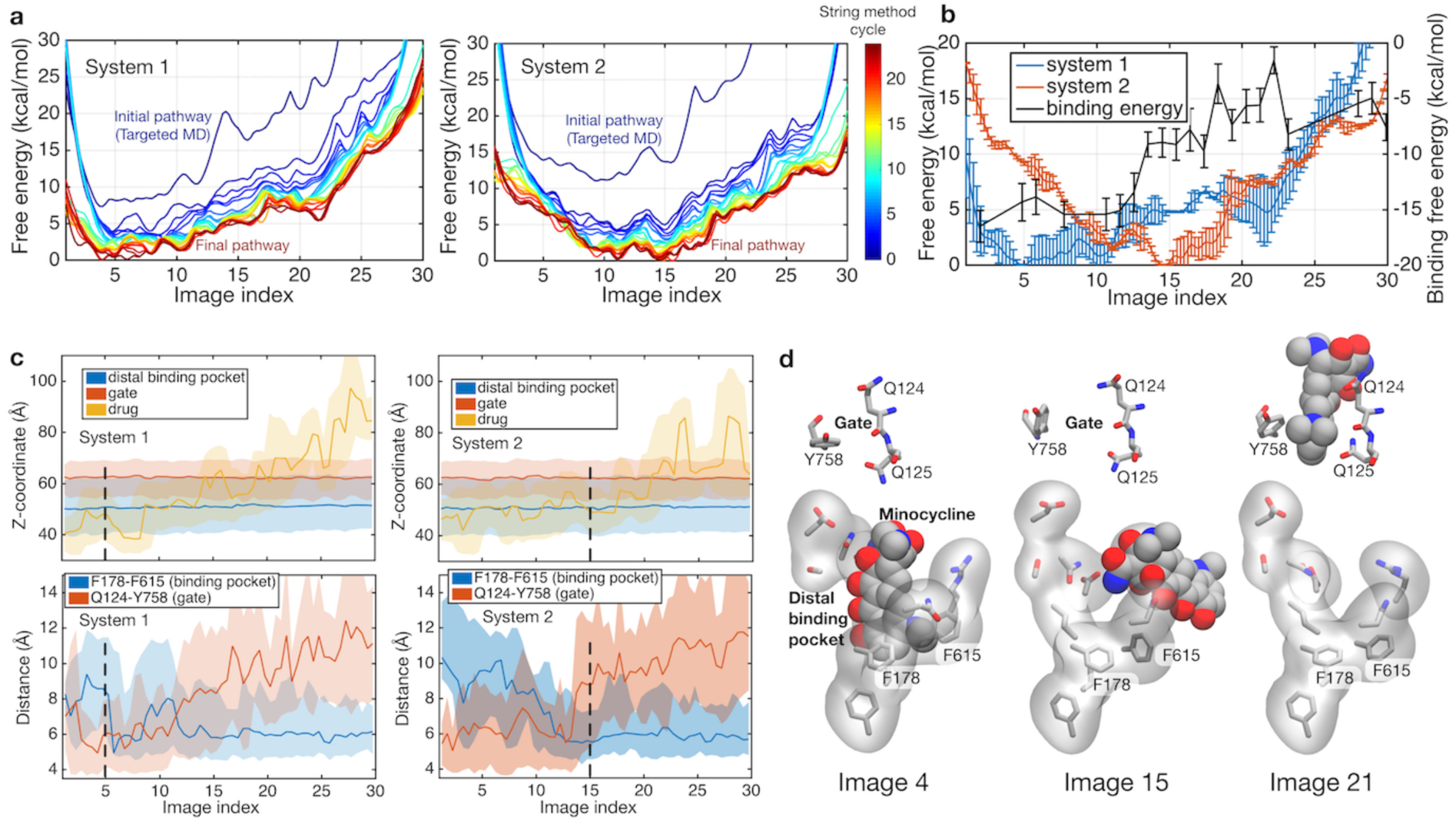
Energetics of functional rotation for two different protonation states. **(a)** Free energy profiles along the conformational pathways obtained by the string method for system 1 and 2. Image 1 represents the BEA state, and image 30 corresponds to the EAB state (see text for the three letter notation). Line color represents the cycle of the string method calculation. The initial pathway (obtained by the targeted MD) is indicated by dark blue, and the final converged pathway is in dark brown. **(b)** Comparison of the final converged pathways for system 1 (blue) and 2 (red). Free energies are referenced to their minima. The black line indicates the absolute binding free energy of the drug for each image of system 2. Statistical uncertainties are represented by error bars. **(c)** *Z*-coordinate (perpendicular to the membrane) of binding pocket residues, gate residues, and drug atoms, and distances between F178-F615 and between Q124-Y758 are shown. Light color regions represent minimum and maximum ranges, and dark lines the average. The positions of minima in system 1 and 2 are indicated by black broken lines. **(d)** Representative structures of the binding pocket and gate residues (sticks), and the drug (spheres) are shown at images 4, 15 and 21.

We further quantified the contribution from the drug-protein interactions at each image by calculating the absolute drug binding free energy to AcrB (see Methods). The absolute binding free energy in system 2 is plotted in Fig 2b with the black line. The binding free energy increases rapidly at around images 10-20, prompting detailed structural analyses for the implication (Figs. 2c and 2d). The closing of the binding pocket was monitored by the distance between the side chains of F178 and F615^31^, and the exit gate opening by the Q124-Y758 distance^14^, as well as the position of the drug (Fig. 2c). During images 5-15, the binding pocket collapses and the exit gate opens, indicating that the minimum free energy state (image 15 in system 2) is ready to release the drug. In system 2, the initial loss of the binding free energy is compensated by the impact of the protonation on the functional rotation (as discussed below). During images 10-20, the drug diffuses into the gate weakly interacting with the tunnel (Fig. 2c). As discussed by Schulz et al^14^ in their targeted MD simulation study, this process may be diffusion-limited. In summary, system 2 represents the protonation state promoting the drug extrusion by the functional rotation.

### Protonation induces a vertical shear motion in the TM domain

How does the protonation at D408 induce the conformational change in the TM domain? To address this question, we first investigated the impact of the protonation on the minimum free energy state (image 5) of system 1 by alchemically transforming it towards system 2, i.e., protonating D408 of protomer I and deprotonating D408p of protomer II (see Methods). Electrostatic potential energy maps before and after the transformation of the protonation state reveal that the TM domain of protomer I is destabilized by the protonation due to a strong positive electrostatic potential bias (Figs. 3a and 3b). This bias is caused surrounding positively charged residues, R971 and K940, whose side-chains orient towards D408p. The relaxation of the potential bias can be seen when inspecting image 15 of system 2, the minimum free energy state (Fig. 3c and Supplementary Fig. S2). The side-chain of R971 (on TM11) reorients downwards (towards the cytoplasm), as does D408p (on TM4) due to repulsion from K940 (on TM10). These motions involve the downshift of TM4 and 11 while TM10 remains at the original position. In order to quantify this relaxation, free energy differences over the alchemical transformation (from system 1 to system 2) were evaluated at images 5 and 15 (see Methods), which yielded 25.9 ± 0.5 kcal/mol at image 5, corroborating electrostatic destabilization of the protonation state in system 2, and only −0.4 ± 6.8 kcal/mol at image 15, suggesting that the protonation no longer affects the energetics at this stage.

**Figure 3:**
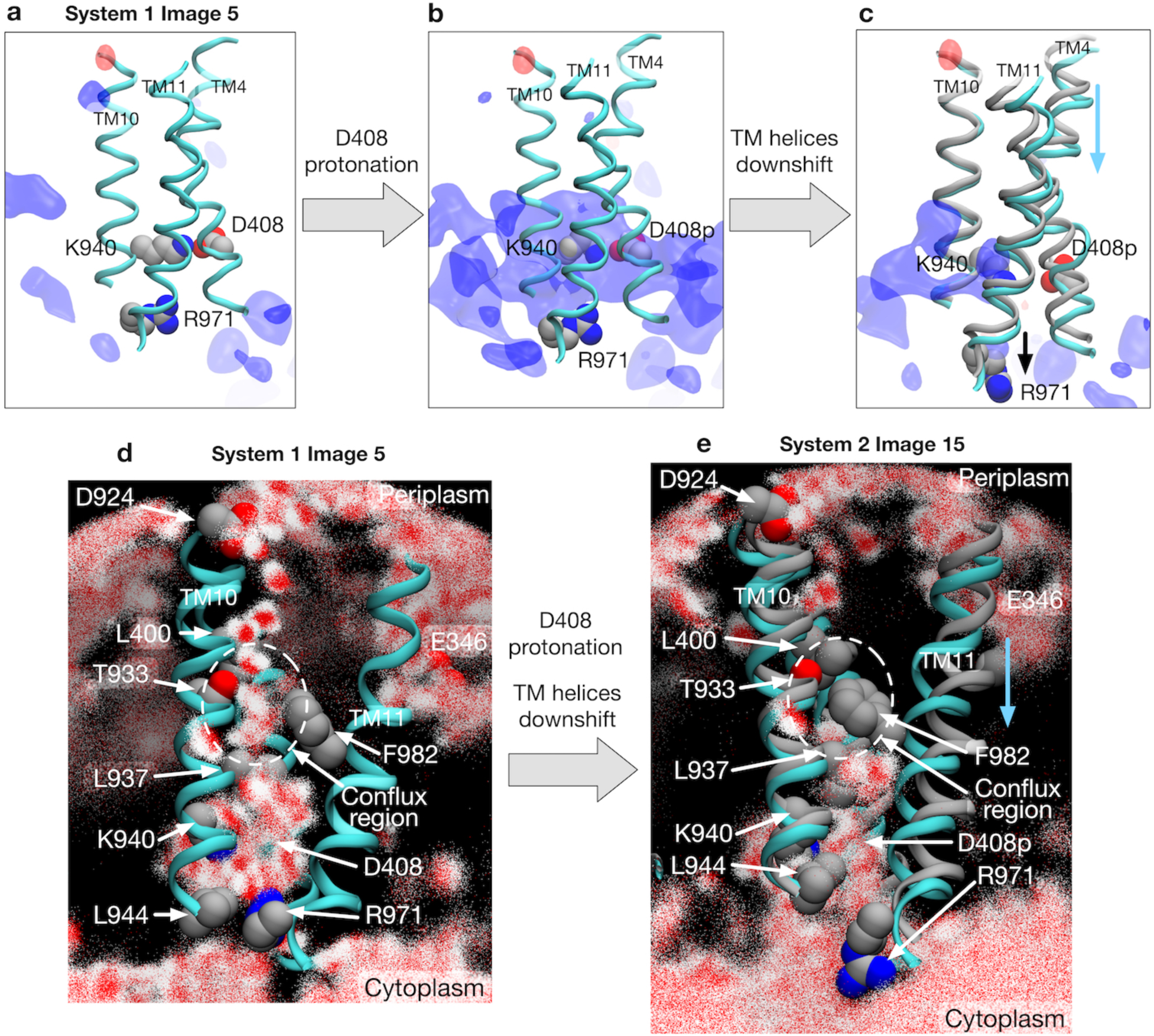
Electrostatic features and water molecule distributions in the transmembrane region. **(a)** Representative structure of transmembrane helices 4, 10 and 11 for protomer I (cyan) of system 1 at image 5 drawn with cartoons. D408(p), K940 and R971 are represented by spheres. Averaged electrostatic potential isosurfaces are drawn in blue (corresponding to the isovalue of 0.06 kcal/mol). **(b)** Just after transforming the protonation state toward system 2. **(c)** System 2 at image 15. For comparison, the helices of image 5 are drawn in gray. **(d)** 2,500 snapshots of water atoms are drawn with red points (oxygen) and white points (hydrogen) for protomer I of system 1 at image 5. Key residues (L400, D924, T933, L937, K940, F982 and R971) are drawn with spheres. **(e)** System 2 at image 15. For comparison, the helices of image 5 are drawn in gray.

By monitoring the centers-of-mass for the other TM helices along the conformational pathway, we found that downshifts occur not only in TM4 and 11 but also in TM3, 5 and 6 (Supplementary Fig. S3). Eicher et al., from the analysis of crystal structures^20^, recognized two rigid assemblies in the TM domain, R1 (TM1 and 3 to 6) and R2 (TM7 and 9 to 12), which changes their mutual arrangement during the transitions among B, E and A states. In terms of R1/R2 rigid body motions, the downshift motions of the TM helices can be interpreted as a vertical shear motion of R1 against R2. Tight packing of the helices within R1 (TM4 and the others), and a rather weak interactions on the interface between R1 and R2 may facilitate such a large rearrangement. TM11 in R2 is the exception, because it moves downwards together with R1. Previous structural studies mostly focused on lateral shear motions and rocking motions^2,20^ of R1 and R2, whereas our observations clarify that protonation mainly induces a vertical shear motion. Indeed, downshifts of the specific TM helices, including TM11, can be discerned in the crystal structures when the TM helices of different states are properly aligned (Supplementary Fig. S4).

### Vertical shear motion in TM domain regulates the alternate access of water

The next question is how D408 become protonated. Because proton translocation across the TM domain requires access of water molecules to supply protons, we investigated the distribution of water molecules in protomer 1 (Figs. 3d and 3e, and Supplementary Fig. S5 for other protomers). At image 5 of system 1 with D408, a water wire channel is clearly observed from periplasmic D924 (a possible transient proton binding site to facilitate proton uptake on TM10)^20^ to the space in the middle of the helices (called the conflux space^32^) surrounded by hydrophobic residues (Fig. 3d). Also, a minor water pathway comes from E346 (another possible transient proton binding site on TM2), which merges into the conflux space. At the cytoplasmic end, the side chain of R971 breaks the channel, preventing the leakage of protons to the cytoplasm.

In contrast, water distribution at image 15 of system 2 with D408p is completely different due to the vertical shear motion in the TM domain (Fig. 3e). There is no water channel coming from the periplasm, due to the collapse of the conflux space by the downshifts of TM4 and 11; distances between residue pairs L400 and T933, L937 and F982 decrease preventing water molecules from entering the conflux space (Fig. 3e). On the other hand, the side chain of R971 opens towards cytoplasmic bulk water, exposing the protonation site and providing a possible proton release pathway. Thus, R971 is an electrostatic gate for proton diffusion to the cytoplasm, analogous to the selectivity filter of aquaporins as discussed in previous studies^20,33^. Mutagenesis studies showed that R971 is an essential residue for proton translocation^34^. In summary, these two states, image 5 of system 1 and image 15 of system 2, exemplify the alternate access of water occurring concurrently with the proton translocation.

### Vertical shear motion in TM domain is coupled to porter domain motion

Next, we studied energy transduction from the TM domain to the porter domain. Does D408p in the B state induce the conformational change of the porter domain? Figure 4 shows the conformational pathways of system 1 and 2 monitored by the distances between D408 and K940 in the TM domain (see Fig. 1d for crystal structure), and between PC1 and PC2 subdomains in the porter domain (see Fig. 1b). The dark blue lines indicate the initial pathways generated by the targeted MD simulations. As shown in previous studies, targeted MD simulation tends to yield pathways where local (TM domain) and global (porter domain) motions are decorrelated with each other^35^. Remarkably, however, the string method yielded synchronized pathways for system 2, but not for system 1. This suggests the existence of a coupling between the remote sites in system 2.

**Figure 4:**
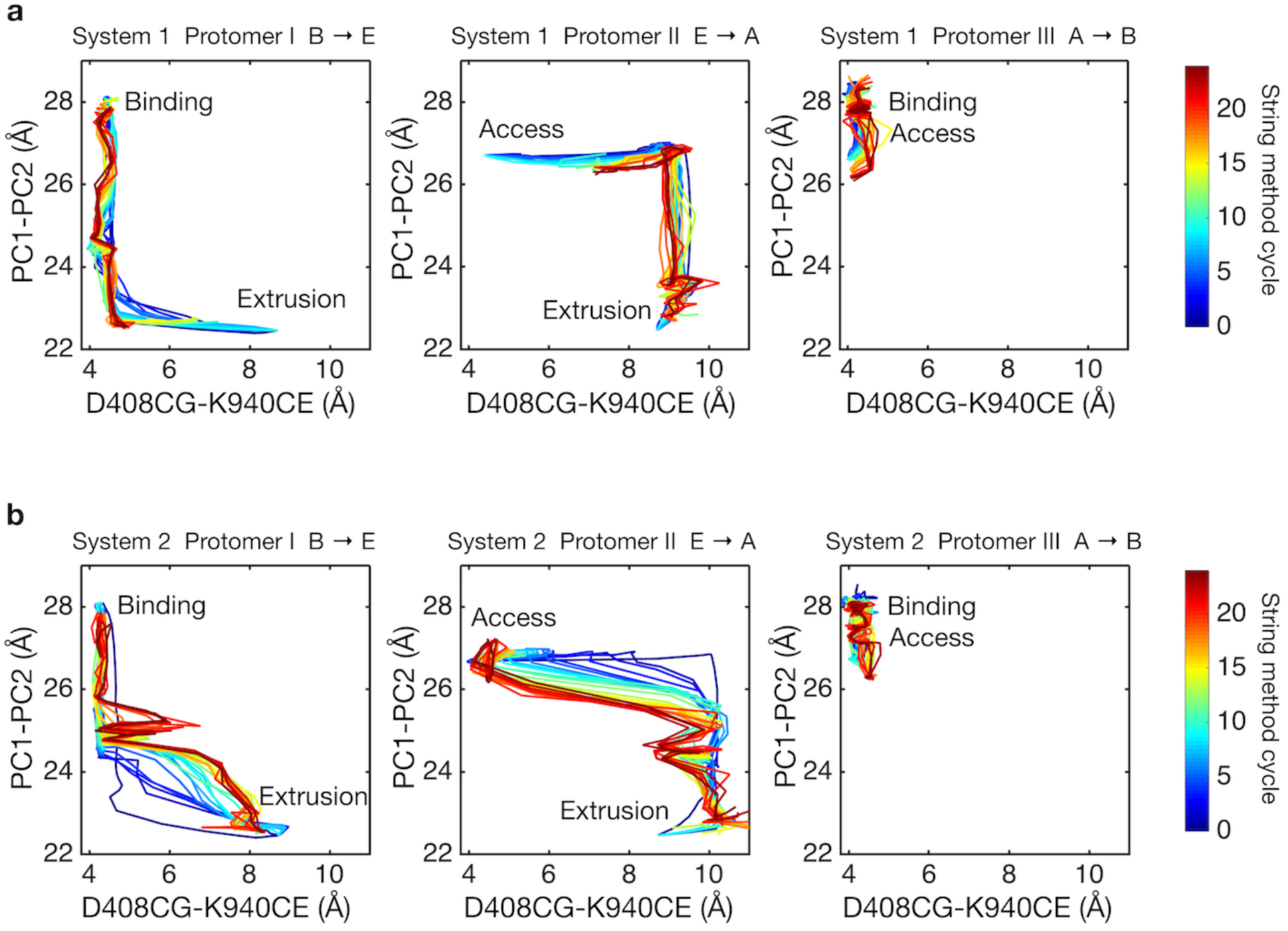
Two-dimensional visualization of conformational pathways. **(a)** Conformational pathways of system 1 are monitored by distance between Cγ-atom of D408 and Cε-atom of K940 (illustrated in Fig. 1d), and the center-of-mass distance between PC1 and PC2 subdomains. These quantities are deterministic because they are involved in collective variables used with the string method calculation. Dark blue lines indicate the initial pathway generated by the targeted MD simulations. Dark brown lines are final converged pathways obtained by the string method. **(b)** Conformational pathways of system 2.

How are the two sites coupled? To quantify dynamic correlations of residue pairs, we employed the mutual information analysis, which has been successfully used in the analyses of allosteric coupling compared with experiments^36^. Mutual information was calculated from snapshots along the minimum free energy pathways obtained with the string method. Figure 5a plots the number of residue pairs between the TM helices and the porter domain as a function of “correlation” measured by mutual information. Clearly, system 2 (broken lines in Fig. 5a) exhibits larger dynamic correlations compared with system 1 (solid lines). Protomer I, among the three protomers, shows largest correlation, indicating that the TM domain motion of protomer I induced by proton translocation is harnessed for the rearrangement of the porter domain. This implies that the transition from the B to the E state is the energy-consuming step^4^.

**Figure 5:**
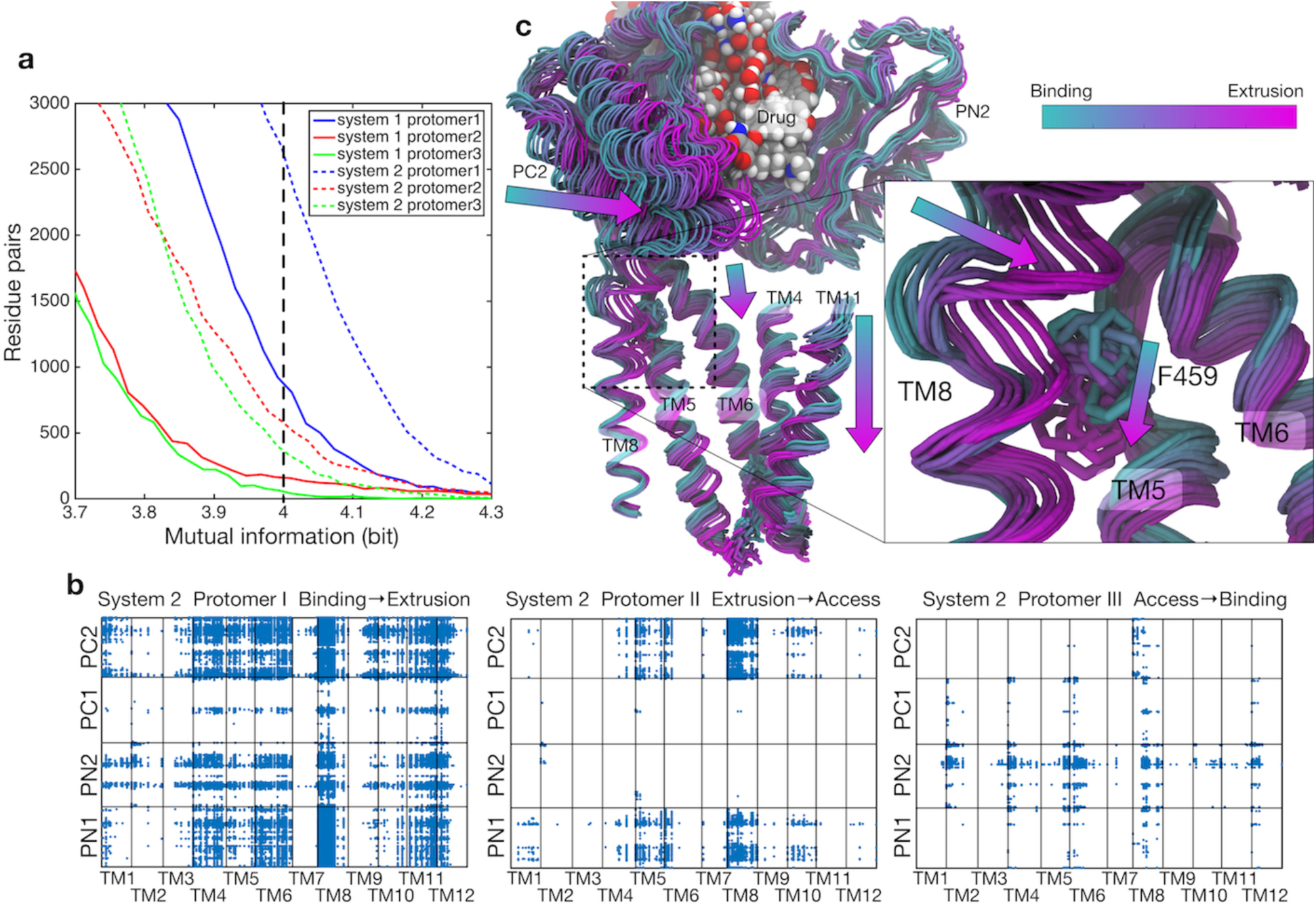
Dynamic correlations between the TM helices and the porter domain motions. **(a)** Number of residue pairs between TM helices and the porter domain as a function of mutual information for each protomer of system 1 and 2. **(b)** Correlated residue pairs between TM helices and the porter domain evaluated by mutual information along conformational pathway of system 2. Each blue dot represents a highly correlated residue pair with mutual information greater than 4.0 bit (indicated by the black broken line in the panel **(a)**). **(c)** Representative snapshots of protomer I along the conformational pathway of system 2. PN2 and PC2 on the porter domain, and TM4-6, TM8 and TM11 on the TM domain, and the drug are shown. According to the image index, the color of the structure changes from cyan (corresponding to the B state) to magenta (the E state). F459 on TM5 and the drug are represented by sticks and spheres, respectively.

Dynamically correlated residue pairs of system 2 are visualized in Fig. 5b (Supplementary Fig. S6 for both systems). Here, residue pairs whose mutual information is larger than a threshold are plotted. In protomer I (B**→**E), high correlations were observed between TM4-6, 8 and 11, and PN1 and PC2 (and also weakly with PN2). As identified above, these TM helices (except for TM8) are involved in the downshift motions induced by D408p. To see how the TM helices and the porter domain are correlated, we visualized the structures along the pathway (Fig. 5c). In particular, the coil-to-helix transition of TM8, which leverages the rigid body motion in PN1/PC2 (see Fig. 1b and Supplementary Fig. S7a), is sterically hindered by the sidechain of F459 on TM5 in the B state (Fig. 5c and Supplementary Fig. S7b). Downshift of TM5 causes the F459 sidechain to flip, allowing the coil-to-helix transition of TM8. Thus, the downshift of TM5 reduces the barrier for the extension of TM8, expediting the PN1/PC2 rigid body motion to open the exit gate for the drug.

Protomer I also exhibits weak linkage between PN2 and several TM helices. These weak correlations may arise from PN2’s rearrangements relative to PC1 due to the shrinkage of the distal binding pocket (see Fig. 1b for the binding pocket location). The downshift of TM helices decreases contacts between the TM domain and PN2 and may accommodate the rearrangement of PN2. Previous studies indicated that a slight tilting of TM2 is coupled to PN2 motion^2,20^, which is also observed here as weak correlations between TM2 and PN2.

Contrary to protomer I, protomer II (E**→**A) has only a local correlation network between TM5 and 8 (and weakly with TM10), and PN1 and PC2, which is associated with the reverse helix-to-coil transition of TM8, closing the drug exit gate. Also, weak correlations can be seen in TM1, 4, and 10, which are consistent with the scenario by Yamane et al^19^ for the E**→**A transition; Twisting motions of TM4 and 10 interfere with the TM1, PN1, and PN2. Protomer III (A**→**B) has rather weak correlations between PN2 and TM helices, which may be related to the rearrangement of PN2/PC1 on distal binding pocket formation and TM helices rising, contacting the PN2 domain. These local and weak correlations in protomers II and III suggest that the minor coupling either occurs indirectly via interfacial interactions between protomers or is mediated by drug binding, not directly from proton translocation.

## Discussion

According to our results as well as the previous studies^11-13,19,20^, we propose the cycle of the functional rotation summarized in Fig. 6. Coupling of the TM and the porter domains during B**→**E is the energy-consuming process. Protonation of D408 in the B state drives a vertical shear motion in the TM domain which regulates the alternate access of water (the conflux space closes, R971 opens towards the cytoplasm) as well as the motion of the porter domain. In this process, the TM8 works as a transmitter. It’s extension induces the rigid body motion of the PN2/PC1 tandem, opening the drug exit gate and shrinking the distal binding pocket. In the E**→**A step, the porter domain of E relaxes towards A without systematic couplings with the TM domain, perhaps mainly driven by cooperativity (interfacial interactions) between protomers. Cross-linking experiments^12,13^ and coarse-grained model simulations^11^ suggested that steric clashes occurred between the E protomers, making two or more E-state protomers in the trimer energetically unfavorable, making the E**→**A transition occur spontaneously. In the A**→**B step, drug bound in A may “catalyzes”^27^ distal binding pocket formation in the PN2/PC1 tandem. The resulting conformational change increases the number of contacts with the TM domain and drives the TM helices upwards.

**Figure 6:**
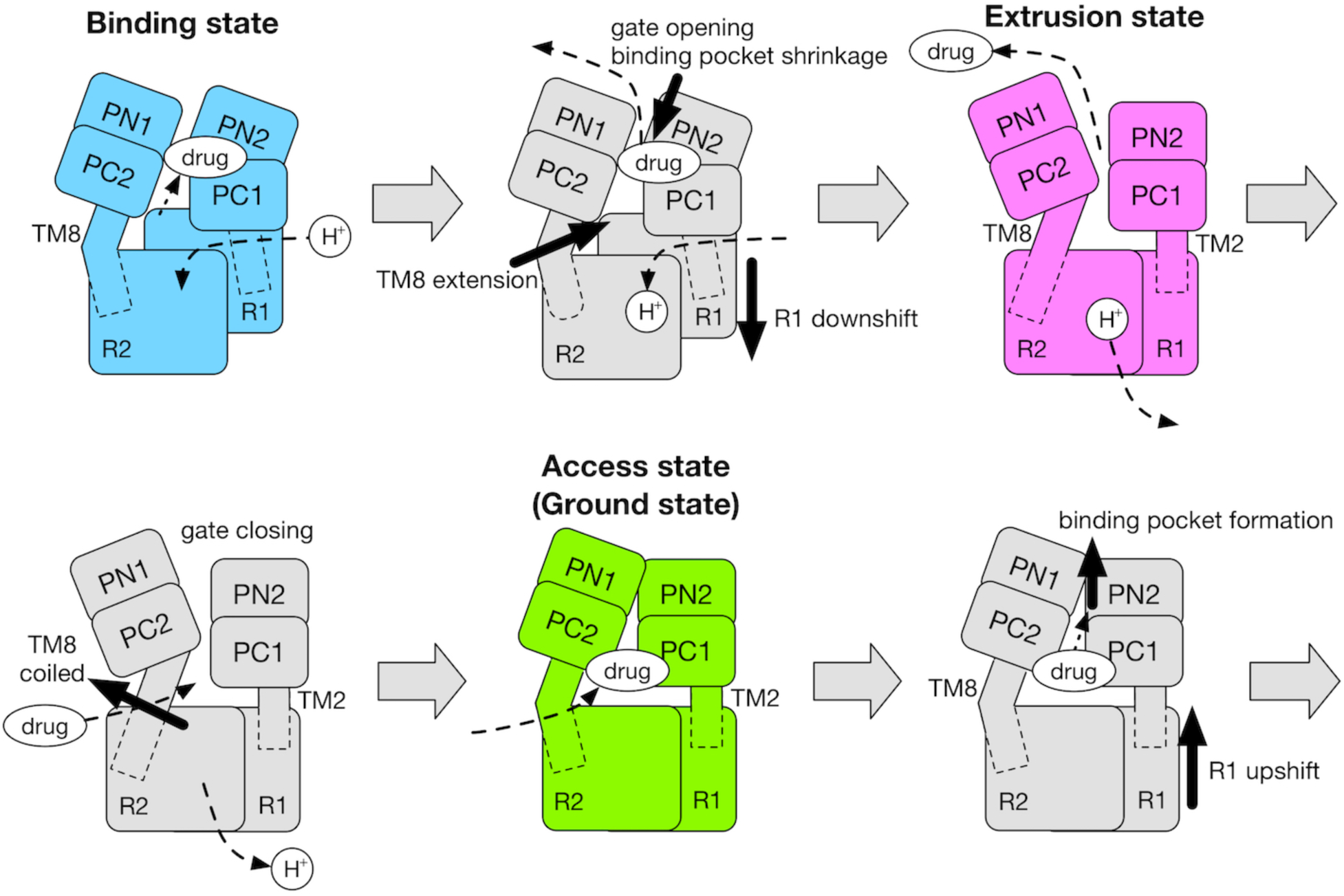
Schematic of the cycle of the functional rotation. Relative motions of the TM domain (consists of R1 and R2 repeats) and the porter domain (consists of PN1, PN2, PC1 and PC2 subdomains) are represented by thick black arrows. The coil-to-helix and reverse helix-to-coil transitions in TM8 were also indicated by thick black arrow. The accessibility of protons/drug to the TM/porter domain is indicated by small black arrows with broken lines. The funnel domain is omitted for visual clarity.

This scheme is consistent with the model proposed by Pos^4^ which is based on a combination of the alternate access model and the binding change mechanism for F_0_F_1_-ATP synthase^37^. The present study provides an evidence that energy is consumed to release drug (i.e., the B-to-E step) in AcrB, which was predicted by Pos’s model in analogy to F_0_F_1_-ATP synthase where the energy is used to release the ATP from the beta-subunit. Moreover, this study provides a simple molecular mechanism for energy transduction from the TM domain to the porter domain in terms of the vertical shear motion in the TM domain. The same type of vertical shear motions can be seen in other transporters which employ the so-called elevator mechanism^38^.

Finally, we discuss the possibility of other protonations which may impact on functional rotation. The computational work by Eicher et al. indicated that not only D408 but also D407 is protonated in the E state suggesting a stoichiometry of two protons per one substrate^20^. On the other hand, pKa calculations by Yamane et al. showed that only D408 should be protonated in the E state^19^. Furthermore, their MD simulations showed that the protonation of both D407 and D408 destabilizes the structure of the TM domain. In this study, we have double-checked our results through the free energy profiles along the pathway and alchemical free energies. The consistency of two independent calculations has confirmed that the impact of transient deprotonation/protonation of D408 is certainly enough to drive functional rotation and extruding a drug.

## Methods

### System setup

The crystal structure of the asymmetric AcrB trimer (PDB entry: 4DX5^31^) with bound minocycline in the distal binding pocket of protomer I (B state) was used for MD simulations. The AcrB trimer was embedded in an equilibrated lipid bilayer membrane of POPE (1-palmitoyl-2-oleoyl-*sn*-glycero-3-phosphoethanolamine). We estimated the TM region using TMHMM^39^ and overlaid it on the equilibrated POPE membrane, then removed lipid molecules within 1.5 Å of the protein. A large hole with a diameter of ~30 Å in the TM center of AcrB should be filled with phospholipids to avoid proton leakage across the membrane. Twelve POPE molecules were manually packed into the hole (six for the upper leaflet and six for the lower leaflet). Then, the simulation box was filled with water molecules. D408 of protomer II (E state) was protonated for system 1 while D408 of protomer I (B state) was protonated for system 2. Finally, sodium ions added to make the net charge of the system neutral. The final system comprised 480,074 atoms.

Equilibration was performed with MARBLE^40^ and mu2lib (publicly available at http://www.mu2lib.org) using CHARMM36 for proteins^41^ and lipids^42^ as force-field parameters. For minocycline, the parameters compatible with CHARMM force-fields, developed by Aleksandrov and Simonson^43^, were used. Electrostatic interactions were calculated with PME^44^, and the Lennard-Jones interactions were treated with the force switch algorithm^45^ (over 8 to 10 Å). The symplectic integrator for rigid bodies^40^ and SHAKE^46^ constraint was used with MARBLE and mu2lib, respectively, using a time step of 2 fs. After an initial energy minimization, the system was gradually heated to 300 K for 1 ns. Then, the system was equilibrated for 8 ns under NPT ensemble (300 K and 1 atm). Throughout the minimization and equilibration, positional harmonics restraints were imposed to the nonhydrogen atoms of AcrB. For the minimization and first 2 ns equilibration, positional restraints were also applied to the nonhydrogen atoms of minocycline and crystallographic water molecules.

### String method

For generating initial pathways to be used in the string method, targeted MD simulations were performed with MARBLE. Starting from the BEA trimeric state, a targeted MD simulation of 10 ns was conducted towards the EAB state. A force constant per atom of 1.0 kcal/Å^2^ was used for the harmonic restraint on the root-mean-square displacement (RMSD) variable measured with the nonhydrogen atoms of AcrB. The target structure (EAB state) was created by rotating the initial structure by 120 degrees around the *z*-axis (perpendicular to the membrane plane). As shown by Schulz et al.^14^, drug efflux towards the exit pore in the targeted MD when pulling only the atoms of AcrB was not observed in 10 ns because the drug extrusion process is diffusion limited. Thus, in order to extrude minocycline toward the exit pore, minocycline was also pulled in our simulation. Starting from the bound form in the distal binding pocket, (the RMSD variable of) minocycline was pulled to 5 Å above the exit gate (Y758) for 10 ns using the same force constant as that of AcrB. The same protocol was applied to both system 1 and 2, yielding very similar trajectories. The trajectories were post-processed for the string method calculation: snapshots were interpolated by piece-wise linear fitting in the collective variable space and 30 equidistant points (called images) were defined as the initial pathway. Image 1 corresponds to the BEA state, and image 30 represents the EAB state. Then, the all-atom coordinates (and corresponding velocities and box size) closest to each image were chosen for the initial structure of the string method.

Collective variables for the string method were carefully chosen according to previous simulation studies. We chose Cartesian coordinates of Cα atoms in the porter domain (PN1, PN2, PC1 and PC2) and selected TM helices (TM4, TM5, TM6, TM8, TM10 and a loop connecting TM5 and TM6) of all protomers (see Supplementary Fig. S1). The chosen porter domain residues were those used in a previous coarse-grained study where it was shown that they successfully capture the essence (thermodynamic and dynamic properties) of drug transportation^11^. The TM helices were chosen because a previous all-atom simulation suggested that those particular TM helices are highly mobile^19^. According to recent studies, which investigated the impact of collective variable choice on pathway accuracy, it is important to choose such mobile domains^47,48^. We also chose Cγ-atoms of D407 and D408 and Cε-atom of K940 as collective variables in order to monitor local side-chain motions (as demonstrated in Fig. 4). The drug was not included in the collective variables since it is diffusive. In total, the collective variables contain 1,659 atoms, consisting of 4,977 Cartesian coordinates.

String method calculations were performed with NAMD^49^ combined with in-house scripts. After defining 30 images along the initial pathway from the targeted MD trajectory, atomistic structure around each atomic structure was relaxed by 35 ns equilibration imposing positional harmonic restraints on the collective variables to its image (Supplementary Fig. S8). After the relaxation, the mean forces type string method^22^ was conducted by using positional harmonic restraints with a force constant of 0.1 kcal/Å^2^. Mean forces were evaluated every 4 ns of simulation and images were updated according to the calculated mean force, then reparameterized to make the images equidistant from each other (also a weak smoothing operation was applied). In order to eliminate external components (translations and rotations) in the mean forces, Cα atoms of TM helices in snapshots were fitted to the reference structure^50^ before mean force estimation. The reference structure was created by averaging the crystal structures of BEA, EAB (120 degrees rotation about the *z*-axis from BEA) and ABE states (240 degrees rotation), thus resulting in a 3-fold symmetric structure^23^. The least-squares fittings of structures were performed only in the *xy*-plane and *z*-coordinates were kept the same as in the original snapshots because membrane proteins are symmetric only in the *xy*-plane. The terminal images (images 1 and 30) were kept fixed during the first 36 ns (9 cycles in the string method), and then allowed to move to relax subtle frustrations in the crystal structure possibly due to detergent molecules or crystal packing. In order to avoid any drifts of the terminal images along the pathway, tangential components to the pathway were eliminated from the mean force when updating terminal images^23^. Convergences of the images were monitored by RMSDs of images from those of the initial pathway (Supplementary Fig. S9). Also, we confirmed that terminal images converged to the same regions as those sampled by brute-force simulations (Supplementary Fig. S10).

In order to obtain more statistics and evaluate free energy profiles along pathways, umbrella samplings were carried out using a single umbrella window per image. Umbrella samplings, each of 30 ns length, were performed around the images of the initial pathway, then over 30 ns around the images after the 9th cycle in the string method, over 10 ns after the 14th cycle, over 15 ns after the 19th cycle, and over 100 ns after the 24th cycle (the final converged pathway). The positional harmonic restraints with a rather weak force constant of 0.01 kcal/Å^2^ were used to achieve sufficient phase space overlaps between adjacent systems. Trajectory data were post-processed by MBAR^24,51^ and weights for the restraint-free condition were obtained. Free energy profiles along the pathways were evaluated by using the progress coordinate^48,52^ (which measures a tangential component) for pathways. To focus on the samples within a reactive tube^53^, distance metric^52^ (which measures an orthogonal component) from pathway was introduced, and samples that deviated further than a cutoff radius of 2,000 Å^2^ were ignored. Changing the cutoff radius to 1,500 or 3,000 Å^2^ did not alter the results qualitatively. Statistical uncertainties (standard errors) in the free energy profiles were estimated from block averages. For analysis of the converged pathway (Figs. 2b, 2c, 5), only samples from those images were used in order to eliminate any biases arising from the pathways of earlier cycles.

For the simulations with NAMD under NPT ensemble (300 K and 1 atm), Langevin dynamics and Nose-Hoover Langevin piston^54,55^ were used. Long-range electrostatic interactions were calculated using PME with a grid size of < 1 Å. The Lennard-Jones interactions were treated with the force switch algorithm (over 8 to 10 Å). SHAKE^46^ and SETTLE^56^ were applied with a time step of 2 fs. A multiple time-stepping integration scheme was used with a quintic polynomial splitting function for defining local force components. Calculations were performed by a hybrid OpenMP/MPI scheme on the K computer of RIKEN AICS^21^.

### Alchemical free energy calculations

In order to quantify contributions from the drug-protein interactions to the free energy profile along the pathway, we conducted alchemical free energy calculations^57^ (double-decoupling method) with NAMD and evaluated the absolute drug binding energy for each image of system 2. The last snapshots of the minimum free energy pathway of system 2 were used for the initial structure. In the calculation, minocycline was gradually decoupled with the other atoms by using a coupling parameter λ with 25 intermediate λ-states (stratifications). The simulation length per each λ-state was 1 ns. During the decoupling simulations, the same positional harmonic restraints (force constant of 0.1 kcal/Å^2^) as the string method were imposed on the collective variables. No restraints were imposed on minocycline because it was hard to define binding sites for images after the shrinkage of the distal binding site. Free energy differences over the decoupling were evaluated by the exponential averages^58^ of the differences of the potential energies between adjacent λ states. Statistical uncertainties were estimated from block averages. The same type of decoupling simulation was performed for minocycline solvated by TIP3P water molecules. The water box size comparable to the AcrB system was used to cancel out the system size effect^59^. From two free energy differences, an absolute binding energy was calculated for each image over a thermodynamic cycle. The error propagation rule was applied to obtain final statistical uncertainties. To reduce computation time, absolute binding energies were calculated only for images 1, 4, 5, 7, 10-23 and 28-30.

Also, to evaluate the impact of protonation of D408 in either the B state or E state, we conducted another type of alchemical free energy calculation. Imposing positional harmonic restraints (force constant of 0.1 kcal/Å^2^) on the collective variables of image 5 of system 1 (D408p in protomer I, and D408 in protomer II), the protonation states were alchemically transformed to that of system 2 (D408 in protomer I, and D408p in protomer II). Because of the restraints on the collective variables, this calculation mainly evaluates the contribution of protonation to the free energy differences. Transformations were carried out in both forward and backward directions, with 36 intermediate λ-states for each direction, resulting in the total simulation length of 58 ns. During the transformation, the total charge of the system was kept neutral. Free energy difference of the alchemical transformation was evaluated by BAR^60^ (implemented in the ParseFEP toolkit^61^) and statistical uncertainty was estimated analytically. The same type of alchemical transformation was also conducted starting from the collective variables of image 15 of system 2 to the protonation state of system 1.

In order to obtain more statistics before and after the alchemical transformation, we conducted MD simulations of 50 ns length for the initial and final state imposing the same type of positional harmonic restraints as above. Then, for the investigation of electrostatic potential energy maps, snapshots taken from these simulations were analyzed by the k-space Gaussian split Ewald method^62^ and instantaneous electrostatic potentials of the smooth part were interpolated and averaged over snapshots on the fixed 3D grids. Most of the analyses were performed with in-house MATAB scripts (publicly available at https://github.com/ymatsunaga/mdtoolbox) and MDTraj^63^. Molecular figures were generated with VMD^64^.

### Mutual information analysis

While the minimum free energy path obtained by the string method provides us with the physically most probable pathway in the collective variable space, it is not useful to characterize
dynamic properties of conformational change at atomistic level (e.g., atoms not involved in the collective variables). Here, in order to characterize the dynamic properties of atomistic fluctuations, we generated reactive trajectories from the umbrella sampling data around the minimum free energy path. The reactive trajectories are defined as the portions of trajectories when, after leaving the initial BEA state, it enters first the final EBA state before returning to the BEA state^65^. For generating the reactive trajectories, we used the property that the minimum free energy pathway is orthogonal to the isocommittor surface^22^. It is known that the distribution of the reactive trajectories follows the equilibrium distribution restricted on the isocommittor surface^53^. Using this property, we resampled the reactive trajectories from the umbrella sampling data according to the weights calculated by the MBAR in the previous analysis. Under discretization by 29 slices orthogonal to the pathway, 100 reactive trajectories were resampled from the umbrella sampling data. Then, mutual information of reactive trajectories was computed for pairs of residues, one from the TM and the other from the porter domains by,

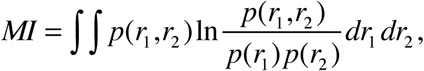

where Cartesian coordinates of the center-of-masses of sidechains were used for *r*_1_ and *r*_2_ because they better capture correlated motions involving semi-rigid regions^66,67^ than the dihedral angles often used in this kind of analysis^36^. *p*(*r*_1_) and *p*(*r*_1_, *r*_2_) are a probability density of *r*_1_, and a joint probability density of *r*_1_ and *r*_2_, respectively. Unlike usual correlation coefficients, the mutual information contains information about both linear and nonlinear dependences. For numerical evaluation of the mutual information, Kraskov’s algorithm^68^, based on *k*-nearest neighbor distances (without binning), was used because it is practical for multi-dimensional data sets such as Cartesian coordinate data.

### Code availability

MATLAB codes developed and used for the analysis of the simulation data are freely available at https://github.com/ymatsunaga/mdtoolbox.

### Data availability

The data that support the plots in this paper and other findings of this study that are not detailed in the Supplementary Information are available from the corresponding author on request.

## Acknowledgment

We acknowledge help from Tomio Kamada and Naoyuki Miyashita in running NAMD on the K computer. We also would like to thank Yuji Sugita, Chigusa Kobayashi, Jaewoon Jung, Motoshi Kamiya, Yohei Koyama, Shun Sakuraba, Luca Maragliano, and Shoji Takada for beneficial discussions. Computational resources were provided by HOKUSAI GreatWave in RIKEN, RICC (RIKEN Integrated Cluster of Clusters) and the K computer in RIKEN Advanced Institute for Computational Science by the HPCI System Research project (Project ID: hp120027). This research was partly supported by Research and Development of the Next-Generation Integrated Simulation of Living Matter, and KAKENHI Grant Numbers 24770159 (to Y.M.) and 25291036 (to M.I.), JST PRESTO (Grant No. JPMJPR1679) (to Y.M.), the Platform Project for Supporting Drug Discovery and Life Science Research from AMED (to K.M., M.I. and A.K.), Innovative Drug Discovery Infrastructure through Functional Control of Biomolecular Systems, Priority Issue 1 in Post-K Supercomputer Development from MEXT (Project ID: hp150269, hp160223 and hp170255) (to T.Y. and M.I.), and RIKEN Dynamic Structural Biology Project (to M.I.).

## Author Contributions

A.K. directed the project. T.T., K.M., H.F., and M.I. contributed to simulation tools. Y.M., and T.Y. conducted simulations. Y.M., T.Y., and M.I. analyzed the data. Y.M., T.Y., S.M., M.I., and A.K. wrote the manuscript. All authors discussed the results and commented on the manuscript.

